# Discovery and characterization of novel inhibitors against ALS-related SOD1(A4V) aggregation through screening of a chemical library using Differerential Scanning Fluorimetry (DSF)

**DOI:** 10.1101/2023.12.20.572618

**Authors:** Maria Giannakou, Constantinos Vorgias, Dimitris G. Hatzinikolaou

**Affiliations:** Department of Biology, National and Kapodistrian University of Athens, Athens, Greece

## Abstract

Cu/Zn Superoxide Dismutase 1 (SOD1) is a 32-kDa cytosolic dimeric metalloenzyme that neutralizes superoxide anions into harmless oxygen and hydrogen peroxide. Mutations in SOD1 are associated with ALS, a disease causing motor neuron atrophy and subsequent mortality. These mutations exert their harmful effects through a gain of function mechanism, rather than loss of function. Despite extensive research, the specific mechanism causing selective motor neuron death still remains unclear. A defining feature of ALS pathogenesis is protein misfolding and aggregation, evidenced by ubiquitinated protein inclusions containing SOD1 in motor neurons. This work aims to identify compounds countering SOD1(A4V) misfolding and aggregation, potentially aiding ALS treatment. The approach employed is drug repurposing and *in vitro* screening of a 1280 pharmacologically active compounds library, LOPAC®. Using Differential Scanning Fluorimetry Technique (DSF), compounds were tested for their impact on SOD1(A4V) thermal stability. Screening revealed one compound raising protein-ligand T_m_ by 7°C, eight inducing a higher second T_m_, suggesting stabilzation effect, and five reducing T_m_ up to 18°C, suggesting possible interactions or non-specific binding.

## INTRODUCTION

### Amyotrophic lateral sclerosis (ALS)

is an incurable motor neuron (MN) degenerative disorder (Deng et al., 1993; Hardiman et al., 2017; Mehta et al., 2019; Wiedau-Pazos et al., 1996). It is the most common form of MNs disease with adult-onset and the third most common neurodegenerative disease (Renton et al., 2014). 50% of patients survives for 3 or more years post diagnosis, 20% for 5 years or more, and up to 10% surviving for more than 10 years (Foyaca-Sibat & Ibañez-Valdés, 2016). Incidence (ca. 1.9 per 100000/year) and prevalence (ca. 5.2 per 100000) are relatively uniform in Western countries (Wijesekera & Leigh, 2009) whereas incidence is lower in Asia (ca. 0.8 cases 100000) (Hardiman et al., 2017).

Based on whether or not a patient has a family history of the disease, ALS is classified as familial (fALS) or sporadic (sALS) (van Es et al., 2017). **fALS** reported rate ranges from 5%-20% due to the related uncertainty around what constitutes family history, and that sALS may sometimes result from gene variants (Mehta et al., 2019). Cu/Zn Superoxide Dismutase 1 (SOD1) mutations were found to cause fALS in 1993 (Rosen et al., 1993) and SOD1 remains the most extensively investigated ALS-related protein while currently, more than 30 different genes have been linked to the familial form of the disease (McGown & Stopford, 2018; van Es et al., 2017). Familial and sporadic forms of ALS are clinically indistinguishable (Simpson & Al-Chalabi, 2006), conferring one common characteristic: the presence of ubiquitinated skein-like or round cytoplasmic inclusion (inclusion bodies, IBs) (Piao et al., 2006) that are SOD1 immunoreactive (Cleveland & Rothstein, 2001; Matsumoto et al., 1996; Shibata et al., 1994).

### SOD1

is a homodimeric copper (Cu) and zinc (Zn)-containing cytocolic enzyme with one intramolecular disulfide bridge per subunit and its main action is to convert the toxic superoxide radicals (O_2_^−^) into water (H_2_O) and hydrogen peroxide (H_2_O_2_) (Abati et al., 2020; Chantadul et al., 2020; Pokrishevsky et al., 2018); (Bruijn et al., 2004).

SOD1’s enzyme activity, depends on the presence of Cu while the role of Zn is related to its folding and stability (Forman & Fridovich, 1973). SOD1 is able to maintain intact the intra-molecular disulfide bond despite the reducing environment of the cytocol. Finally, two Cu/Zn-loaded and oxidized subunits are held together by means of hydrophobic contacts (Rakhit et al., 2004) that further exhibits a very high dimerization constant (K_d_ ca. 10^−10^ M^−1^). These posttranslational modifications endow the protein with high stability (Furukawa et al., 2006; Srinivasan & Rajasekaran, 2018); they are able to shift its unfolding transition from body temperature (Furukawa et al., 2016) for some ALS mutants up to 75°C for the fully mature WT enzyme (Furukawa et al., 2016; Wright et al., 2016).

### Mutations

distort the native conformation of SOD1 and render the protein prone to lose its PTMs and consequently its characteristic stability, exerting gross effect on monomer-dimer equilibrium (Rakhit et al., 2004). So far more than 190 ALS-related mutations have been identified (https://alsod.ac.uk/) in SOD1 (Figure 1). Most of these mutations result in aberrant folding of SOD1, thereby leading to the formation and accumulation of oligomeric and aggregated species of the protein (Bruijn et al., 2004) and/or altering enzyme’s activity (Crown et al., 2019; Zu et al., 1997) ALS-related mutations of SOD1 are scattered through the entire sequence of the SOD1 protein, suggesting an impact on the whole protein (Banerjee et al., 2016; Sau et al., 2007).

### A4V mutation

accounts for almost 50% of ALS-related SOD1 mutations in the USA but is rare in Europe (Andersen et al., 2003). SOD1(A4V) provokes a rapid and aggressive form of fALS reducing survival time to less than 2 years (Saeed et al., 2009). The A4V mutation destabilizes SOD1 protein in both the apo and metallated states (B. F. Shaw et al., 2006; B. Shaw & Valentine, 2007), involving distortion of the structure of the protein and exposure of the hydrophobic core, destruction of β-barrel structure and the copper binding site, along with severe structural changes of the dimer interface (Kim et al., 2014; Lindberg et al., 2005).

### The exact mechanistic details of how SOD1 aggregates cause toxicity

are still being investigated. However, several consequences of the aggregates include: sequestration of other proteins essential for cellular components (Cleveland & Liu, 2000; Johnston et al., 2000), reduction of the availability of chaperones needed by other proteins (Cleveland & Liu, 2000; Rakhit et al., 2004), disruption of the ubiqiutin-proteasome pathway by impairing its ability to degrade not only the mutant protein itself but also other proteins (Cleveland & Liu, 2000; Rakhit et al., 2004). (Figure 2).

Despite its severity; ALS still remains **incurable** (Mehta et al., 2019). Current drugs available for ALS patients and approved by the US national Food and Drug Administration, are: (i) **Riluzole**, whose mechanism of action is related to the inhibition of pre-synaptic glutamate release (Wijesekera & Leigh, 2009), (ii) **Edaravone,** an antioxidative compound reducing oxidative stress (Mejzini et al., 2019; van Es et al., 2017), (iii) **Tofersen**, an ASO decreasing SOD1 protein synthesis (Miller et al., 2013), (iv) **Sodium Phenylbutrate and taurursodiol**; related to the amelioration of the production of mitochondrial energy (Fels et al., 2022; Kubota et al., 2006) and (v) **Dextromethorphan HBr and quinidine sulfate** as symptomatic treatment for pseudobulbar affect (Smith et al., 2017).

ALS is characterized as a rare disease (<50 cases per 100000 (Shah et al., 2021)), however, it still constitutes significant socio-economical burden in terms of healthcare cost (Musteikyte et al., 2020). Consequently, there is urgent need for discovering effective therapies against ALS. Since protein misfolding and aggregation are considered as central events in ALS pathogenesis, therapeutic strategies targeting these processes hold promise (Elliott et al., 2020). The understanding of SOD1 aggregation kinetics has introduced new possibilities for ALS therapy, by the discovery of small molecules that bind and stabilize either (i) the native state of SOD1 or (ii) the intermediate aggregate species (misfolded monomers, low molecular SOD1 aggregates) (Banerjee et al., 2016; Wang et al., 2003), (Rakhit et al., 2004).

### Pharmacological chaperones

are small-molecular-weight molecules that can bind onto specific proteins and promote proper folding and stability, thereby inhibiting aggregation and/or degradation (Bose & Cho, 2017; Convertino et al., 2016). Since their first description in the early 2000’s (Fan, 2003) pharmacological chaperones have entered the clinical research and practice for some rare conformational diseases (Liguori et al., 2020; Tran et al., 2020) e.g. tafamidis against transthyretin amyloidosis (ATTR), stabilizing the transthyretin (TTR) tetramer (Bulawa et al., 2012; Maurer et al., 2018), migalastat that assists proper folding of Fabry Disease (FD)-related α-galactosidase (GLA) variants (Germain & Fan, 2009), etc.

Drug discovery practice demonstrates that usually only one out of thousands compounds may be a viable drug option. For ALS drug discovery is even more challenging given the inherent difficulties to identify compounds that bind to a mixture of transient aggregation species during aggregation kinetics, the fact that there are no natural ligands of SOD1 to serve as a molecular scaffold (Ray et al., 2005) and also mutations’ heterogenous effects. In this study we choose to identify rescuers of SOD1 misfolding by screening a commercially available library, LOPAC® (<u>L</u>ibrary <u>o</u>f <u>P</u>harmacologically <u>A</u>ctive <u>C</u>ompounds), using calculated T_m_ value, as proxy of protein stability. This library contains 1280 marketed drugs or well-characterized small molecules displaying lead-like or drug-like properties, and to our knowledge this is the first time LOPAC® library is used in the context of a random screening against ALS-related SOD1 variants for the discovery of inhibitors of its aggregation (protein-drug interaction). Thermal shift assay (TSA) or more commonly described as differential scanning fluorometry (DSF) (Pantoliano et al., 2001) was chosen for the screening due to a combination of advantages including high sensitivity, its applicability to be used at medium or high throughput level, the low cost, the low concentration of protein (range of a few μM) and tested compounds needed (mM), and the simple requirement of the RT-qPCR machine commonly available in every lab (Gao et al., 2020; Lo et al., 2004; Niesen et al., 2007).

## MATERIALS & METHODS

All chemical reagents (buffers, solutes, inducers, antibiotics DTT, EDTA, and media were purchased from Merck/Sigma-Aldrich (USA).

ATC (Anhydroteracycline) is used as the inducer for tetracycline-inducible expression system (Baumschlager et al., 2020) that pASK75 vector carries. The stock solution is prepared 2mg/mL ATC and it is dissolved in pure ethanol, light protected and stored at −20°C. The working solution is diluted to 0.02mg/mL=20μg/mL in ethanol. The last solution is added to the cultures diluted 1/100.

All solutions are filtered with 0.2μm syringe filters (Corning, US) and transferred to sterilized vials.

Propagated plasmids were purified using NucleoSpin® Plasmid from Macherey-Nagel, MN (Germany) or Plasmid Midi kits from Qiagen (Germany).

For recombinant h-SOD1 purification from bacterial cells, Ni-NTA Agarose (part of QIA*express* Kits, Qiagen (Germany) was used.

Sypro Orange Protein stain 5000X concentrate in DMSO, (Thermo-Fischer, USA) that was diluted as described for the needs of each experiment using the appropriate buffer or solution.

### *E.coli* cultures & hSOD1 recombinant production

*E.coli* cells BL21 trxB freshly transformed with the expression vector pASK75-flag-SOD1-6His were used for protein production. Single colonies were picked and used to inoculate overnight liquid cultures of 6mL. Cells from overnight cultures diluted to OD_600_ 0.1-0.5 were used to inoculate cultures of 250mL or 500mL LB media containing 100μg/mL ampicillin (pASK75 expression vector) and 50μg/mL kanamycin (BL21 trxB cells’ resistance) as well as 200μΜ CuCl_2_ and 200μΜ ZnSO_4_ (salts carrying the metals/co-factors of SOD1). Cultures were grown at 37°C with adequate aeration in shaking incubator and when OD_600_ reached to 0.7-0.8, SOD1 production was induced by the addition of atc in final concentration of 200ng/mL. Temperature was switched to 19°C for overnight induction of about 19hr.

### SOD1 purification

Cell pellets from 250mL culture were collected after SOD1 overexpression cultures by centrifugation at 6000rpm for 15min at 4°C (Juan KR25i, DJB CareLab, UK). Cells pellet is washed once with 20mL lysis buffer and centrifuged again with the same conditions. Cell pellet can be used straight away or can be stored at −80°C for up to 5 months. They are resuspended in 19mL lysis buffer (50mM NaH_2_PO_4_, 300mM NaCl 10mM imidazole, pH 8.0) and are lysed by mild sonication in ice: 75% power, 1sec on 9sec off for 15-18min in a probe sonicator (Vibra Cell^TM^, Sonics & Materials Inc, US). The soluble cell lysate is collected by means of centrifugation (FastGene HighSpeed Mini Centrifuge, Nippon Genetics Europe) at 13000 rpm for 25min at 4°C. 1 mL of Ni-NTA agarose resin (Qiagen, Germany) is packed onto a propylene chromatography/cartridge column (Qiagen, Germany). Column is washed adequately with water before use and equilibrated with 15CV with lysis buffer. Soluble cell lysate is then loaded onto the equilibrated column and the His-bound protein is washed with 15CV wash buffer (50mM NaH_2_PO_4_, 300mM NaCl 20mM imidazole, pH 8.0). SOD1 is eluted off the column (after 0.5mL of dead volume is discarded) with elution buffer (50mM NaH_2_PO_4_, 300mM NaCl 250mM, imidazole, pH 8.0) in 3 elutions steps of 0.5mL, 1mL and 0.5mL or 2mL in total. For the subsequent SOD1 quantification, UV spectrophotometer (Shimadzu, Japan): molar extinction coefficient at 280 nm is 5500 [1/M·cm] and Abs 0.1% [1g/L]=0.361, (Wright et al., 2016) was used.

### Gel filtration Chromatography

The second elution fraction following the first purification step (Ni-NTA gravity flow chromatography) of SOD1 is further purified by size-exclusion chromatography (SEC) using a custom made gel filtration column (dimensions: id 1.2×100cm) packed with Sephacryl^TM^ S-200 (GE Healthcare, USA) in order to to isolate the dimeric protein fraction in TBS buffer, pH 7.4. The chromatographic system is comprised of a pump ISMATEC (Thermo Fisher, USA) and a UV detector Pharmacia LKB-Uvicord SII (American Laboratory Trading, USA).

### In gel activity measurement

SOD1 enzymatic activity was monitored using the riboflavin/nitroblue tetra-zoleum (NBT) gel-based assay. Ten μL of SOD1 WT or mutant of concentration 30-40μm with or without the selected compounds after being diluted with Laemli Buffer (that did not contain SDS or reducing agent) were loaded onto the wells and then analyzed in a 10% polyacrilamide gel. Following electrophoresis the gel was washed thrice with dH2O for 5min each wash. The gel was stained using 12.5mL of a solution that contained: 50mM Sodium Phosphate buffer pH 8.0, 65.9mg Riboflavin, 3.8mg NBT and 52μL TEMED after incubation for 1hr at 4°C and at dark while shaking. After exposure to light SOD1 activity can be seen as colorless bands on the gel because at the areas where SOD1 was, it scavenged the free radicals inhibiting the reduction of the NBT that gives the blue color by the formation of the insoluble formazan.

### Differential Scanning Fluorimetry (DSF) measurements

DSF was carried out using a Biorad CFX Real-Time PCR machine (USA). Proteins at 80μM for SOD1 WT and 20μm for A4V mutation with or without the selected compounds were mixed incubated for about 15min under mild shaking and were measured thereafter. Final concentration of DMSO in the mixture was 0.02% for the samples during LOPAC® library screening. SYPRO Orange dye (Life Technologies) dye at 10× concentration was used as the florescent probe. Sypro Orange dye (excitation and emission wavelengths: 492 and 610nm, respectively (Sorrell et al., 2010)), was chosen due to its high signal-to-noise ratio (Niesen *et al*. 2007).

For the LOPAC®screening: the compounds were diluted 20 times in TBS buffer from the stock solution with initial concentration 10mM and 10% DMSO resulting in final concentration of 500μM. The freshly purified SOD1(A4V) dimeric protein was then added in a concentration of 20μM, thus making the ration between SOD1(A4V): compound equal to 1:20. The initial mastermix of the SOD1 protein with SYPRO orange was added at each rack containing the compounds of the library and was mildly pippeted. Then a small incubation of 20min at RT followed onto a shaking platform set at minimal rpms, while the rack was light-protected. Each compound was tested in triplicates. Each 96-well contains as control sample SOD1(A4V) and SOD1WT. Temperature scan was performed from 4°C to 91°C at a ramp rate of 2°C/min. The analysis of melting temperature (T_m_) was calculated from the minimum of the first derivative of the RFU curve extracted from the PCR apparatus. Data are presented and analyzed using the online tool: https://gestwickilab.shinyapps.io/dsfworld/.

## RESULTS

### Purification of h-SOD1

Achieving a satisfactory recovery yield in the elution fractions proved to be a difficult task due to SOD1(A4V) misfolding and high aggregation propensity that leads to less accessible His and subsequently much fewer interactions with the nickel beads. As a result, the majority of the protein was lost in the flowthrough fraction and wash fractions.

Optimal results were obtained with milder overexpression conditions and after IMAC purification protocol optimization, a satisfactory yield was achieved (Figure 3, Figure 4).

### Size Exclusion Chromatography

It is important to evaluate the ratio among the different conformational species of SOD1 and separate the dimeric form of the protein since that is the physiological state of the protein and this is the target of the compounds tested in this study. MutSOD1 dimer is less stable and thus it remains lesser time in this state compared to the WT dimer (Ayers et al., 2017; Furukawa et al., 2016; Lindberg et al., 2005).

As seen from Figure 5 and Figure 9, for WT the majority of the species that dominates is the dimeric form while for A4V and and despite the milder possible overexpression conditions, this form is significantly decreased (Figure 7 and Figure 9) For WT there is a small amount of protein (since it is spotted via SDS-PAGE, (Figure 6) at fraction 10 corresponding to insoluble aggregates; possibly to a trimeric species of SOD1 that is negligible in quantity relatively to the dimer and that could not be detected in Native Gel (Figure 9).

For A4V case, it is obvious from the ratio between the dimeric form and the proceeding peaks corresponding to aggregation species, that the dimer state of the protein is noticeably smaller (Figure 7). A considerable portion of the protein is eluted earlier in context of a complex and not separated wide peak (elution fractions 7 to 11) which corresponds to bigger molecular weights as seen from Native-PAGE ((Figure 9). This aggregate species also contain an insoluble component as seen form SDS-PAGE corresponding possibly to a trimeric species (similarly to WT).

### Activity of SOD1 enzyme

The 2^nd^ and 3^rd^ elution fractions derived by IMAC purification of each SOD1 were tested. We chose to test SOD1 activity in the 2^nd^ elution fraction (derived by IMAC purification), due to its higher purity and concentration (Figure 10a). Based on the zymogram assay, both SOD1 WT and A4V are qualitatively active. Quantitative analysis of the band intensity after equimolar loading, using the Image J software, suggests that SOD1(A4V) activity is almost the 87±3% of the SOD1 WT activity (Figure 10b). Furthermore, denaturation at 95°C leads to a 95±1% loss of activity for both variants. Lastly, the presence of an equimolar amount of EDTA chelator reduces by 20% the scavenging capacity of SOD1(A4V) in contrast to the unaffected SOD1 WT activity (Figure 10b, c)

**Figure 10:**
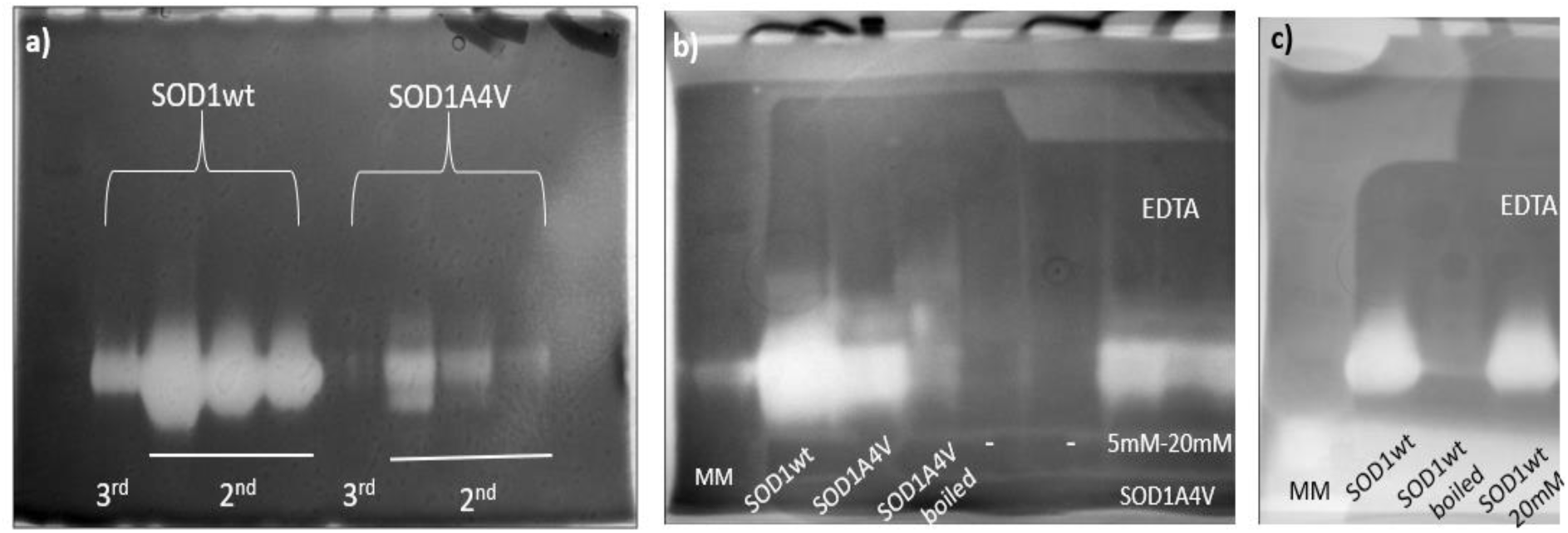
Zymogram performed by the method of riboflavin and B2/NBT. At the first gel a) from the left: SOD1WT: 3^rd^ elution in dilution 1:1, and the following three from the 2^nd^ elution in concentrations 20uM, 8μm and 4uM, while the following four lanes are SODA4V: 3^rd^ elution in dilution 1:1 followed from the 2^nd^ elution in concentrations 13μM, 5μm and 2μM. At the At the second gel b): equimolar quantities of the WT and A4V SOD1 variant, SD1A4V boiled, and SOD1(A4V) incubated 15sec before loading onto the gel with EDTA. At the third gel c) SOD1WT, SOD1WT boiled and SOD1WT incubated 15sec before loading onto the gel with EDTA.

### Selection of the chemical Library: LOPAC®

The library chosen to be screened to find potent rescuers of aggregation for the pathogenic ALS-related mutant is LOPAC® (<u>L</u>ibrary <u>o</u>f <u>P</u>harmacologically <u>A</u>ctive <u>C</u>ompounds). It is a commercially available library containing 1280 marketed drugs or well-characterized small molecules displaying lead-like or drug-like properties. We chose that library for the following reasons. Firstly, it is a widely used library containing compounds displaying multiple mechanisms of action and in addition it covers the field of cell signaling and neurotransmission. Secondly, these compounds target a large number and diverse set of biological receptors. Thirdly, it covers a broad range of molecular weight from 36 to 1485 Da (Vliet et al., 2017). To our knowledge this is the first time LOPAC® library is used in the context of a random screening against ALS-related SOD1 variants for the discovery of inhibitors/rescuers of aggregation that are directly targeted to SOD1 (protein-drug interaction). In another study LOPAC library was assessed in MN cells aiming to discover compounds that decrease neuronal damage in the presence of mutSOD1 (https://www.axionbiosystems.com/sites/default/files/michael_hendrickson_-_brainxell_-_sfn2017.pdf).

### Optimizing the conditions before screening

The desirable buffer for the organic library screening should mimic physiological conditions and thus Tris Buffer Saline (TBS) and Phosphate Buffer Saline (PBS) were examined. Gel Filtration chromatography (SEC) using these buffers was performed and DSF spectra were extracted (Figure 11).

**Figure 11:**
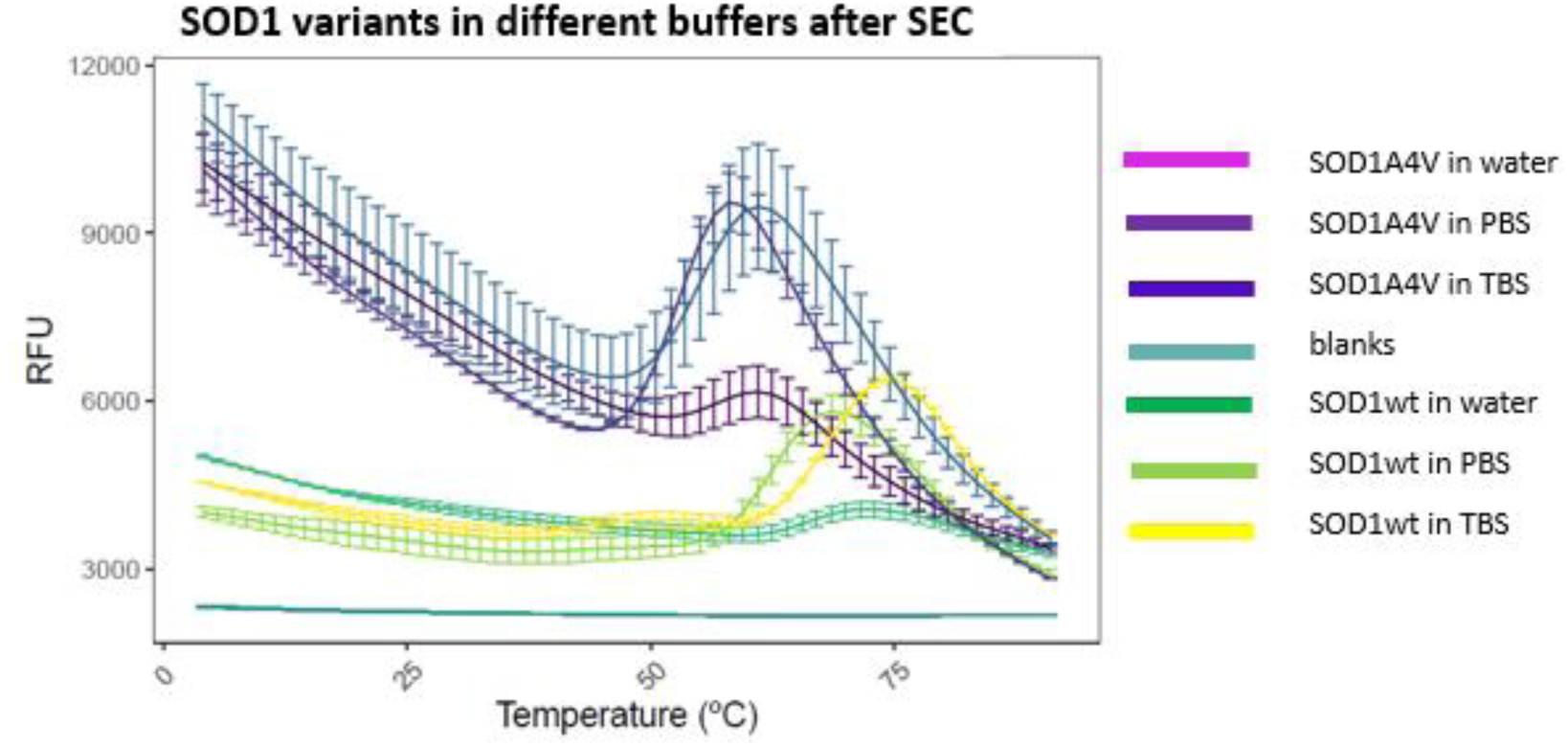
Examination of different buffers in SOD1 stability.

Based on the calculated T_m_ (Table 1), it is obvious that when both SOD1 variants are resuspended in the two physiological buffers tested, a significant increase in T_m_ is observed compared to the initial buffer tested (Table 1, Figure 11) (elution buffer, containing 250mM imidazole and two-fold NaCl concentration compared to TBS, PBS). TBS is chosen for the subsequent screening of the library, because it presents a slightly larger increase in T_m_ compared to PBS buffer, possibly implying that it is the more appropriate for SOD1; nevertheless, the difference in T_m_ attributed by the two buffers is minimal for both SOD1 variants.

**Table 1:** Calculated T_m_ following SEC of both SOD1 variants using different buffers.

| Condition | $T_m$ (°C) | SD |
| --- | --- | --- |
| SOD1(A4V) in water | 56.2 | 0.5 |
| SOD1(A4V) in PBS | 52.3 | 0.1 |
| SOD1(A4V) in TBS | 54.7 | 0.2 |
| SOD1WT in water | 66.1 | 0.1 |
| SOD1WT in PBS | 62.1 | 0.0 |
| SOD1WT in TBS | 67.9 | 0.1 |

### LOPAC® Screening

The main idea (Figure 12) is to incubate the library with SOD1(A4V) for an adequate amount of time to allow for interaction and to monitor for T_m_ shifts (if any). In general, higher T_m_ values correlate with a more stable protein (Niesen et al., 2007; Pantoliano et al., 2001). If the candidate drug molecule added in the mixture binds specifically to the protein, it can stabilize the protein and cause an increase in T_m_. An increase in T_m_ of 1-2°C can be considered as a sign of protein-drug interaction (Huynh & Partch, 2015; McMahon et al., 2013).

**Figure 12:**
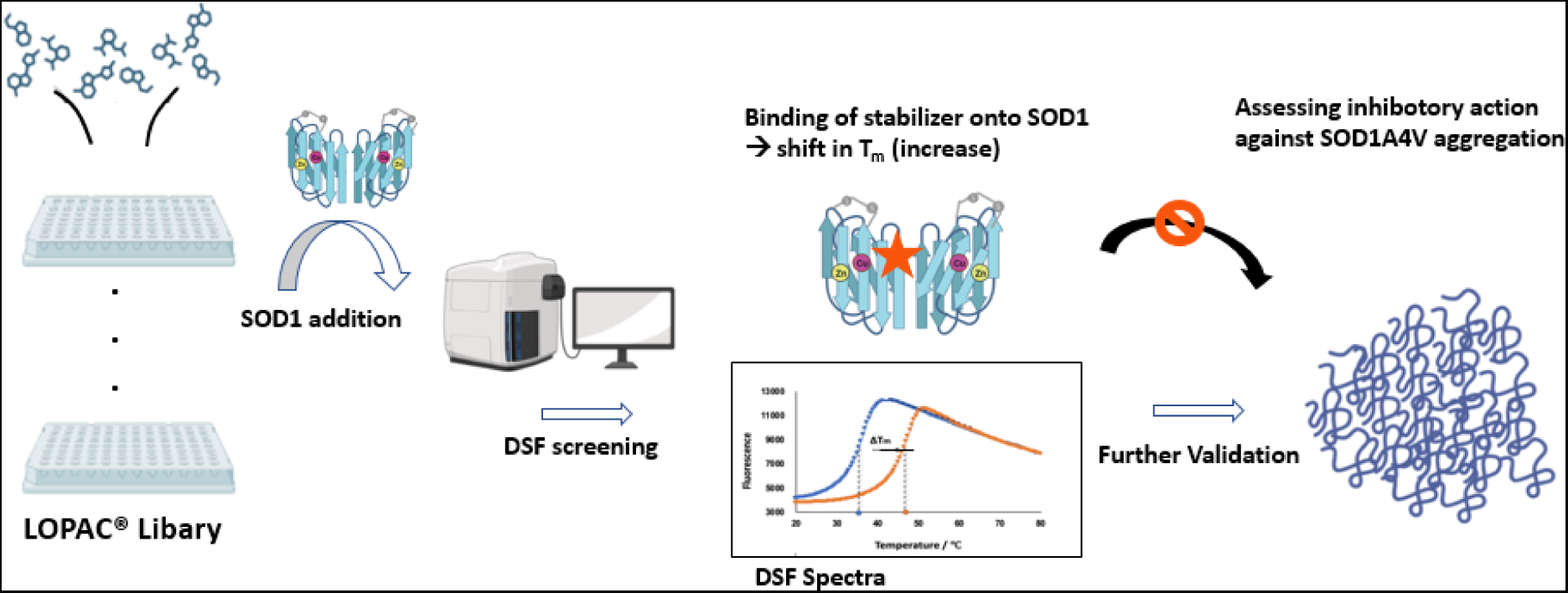
Workflow of LOPAC® screeing. The compounds are arranged (after proper dilution) in a 96-well plate format and the freshly purifies SOD1(A4V) is added to the mixture. Following a short incubation DSF are extracted. The compounds presenting a positive shift in the T_m_ of the mixture will further be evaluated as potent aggregation inhibitors of SOD1 in kinetics experiments. (Created in BioRender.com).

### LOPAC® Screening Results

Interestingly the majority of the LOPAC® compounds induced a difference in the T_m_ of the protein towards both directions: increasing it and reducing it. All ΔT_m_ values are presented at Supplemental Table 1.

To have an overview of the effect of LOPAC® results the following figure (Figure 13) depicts quantitatively the difference in SOD1’s T_m_ caused by the presence of each compound.

**Figure 13:**
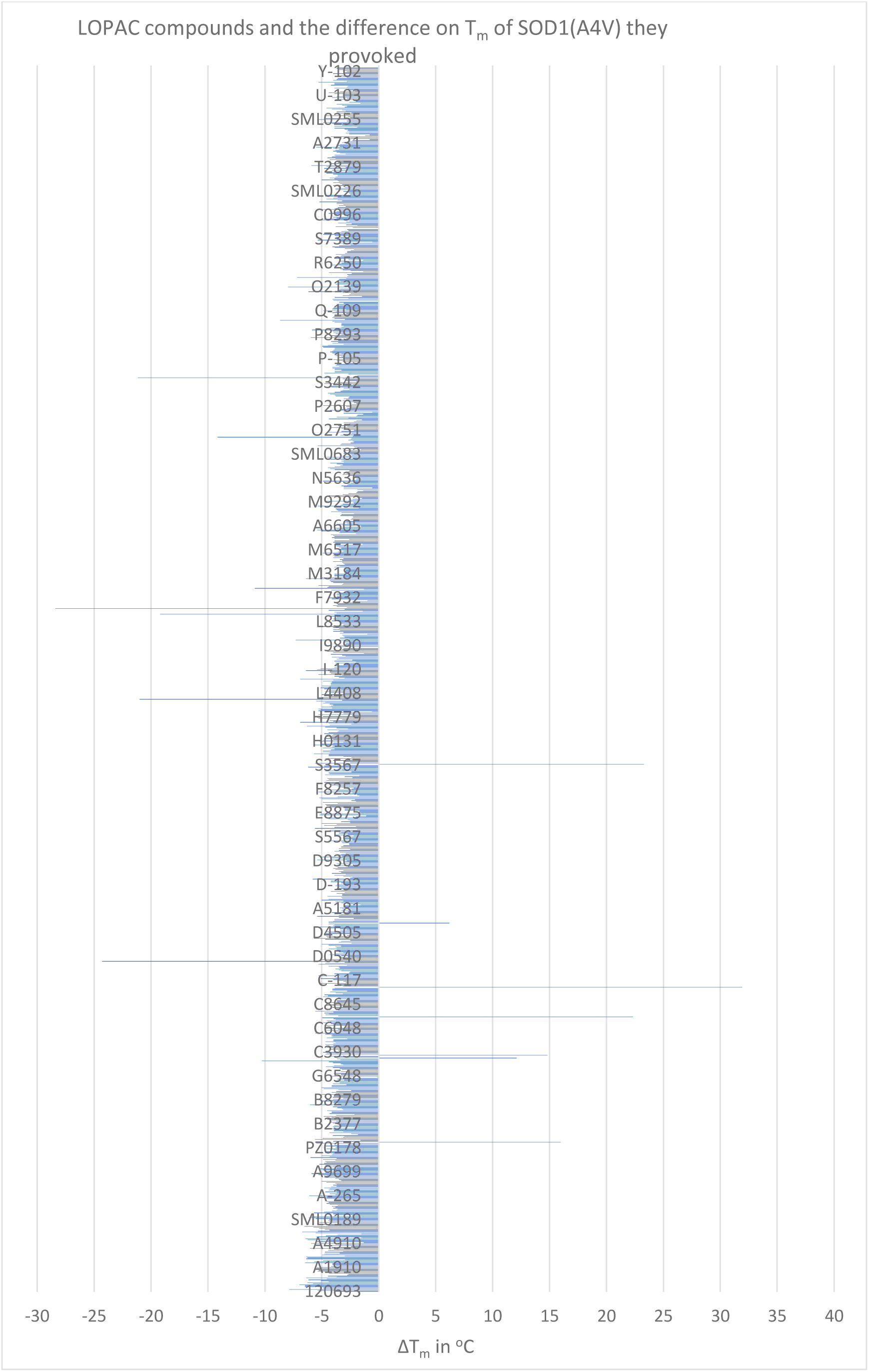
The difference on T_m_ of SOD1(A4V) caused by the compounds of LOPAC® library. ΔT_m_ corresponds to the function: T_m_(SOD1(A4V))-T_m_(SOD1(A4V) upon compound’s presence).

Almost all compounds or 99.5% of the total number of 1280 LOPAC®’s compounds (except for 7), presented a negative ΔT_m_, or in other words they were able to decrease the T_m_ of the protein from 6 to almost 30°C. From the bulk of the compounds that presented a negative shift, 1260 of them corresponding to the 98.4% of the total number of the library chemicals, induced a negative shift in SOD1(A4V) T_m_ resulting in a difference from −7°C to −0.5 °C. However, 0.5 °C difference is small and generally considered negligible in assessment of protein binders, where a difference >1°C is considered as interaction (Huynh & Partch, 2015; McMahon et al., 2013).

Following the analysis of LOPAC® screen results, the distribution pattern of the algebraic value of the ΔT_m_ is correlated with the number of compounds that provoked it, separated in certain ranges (Figure 14).

**Figure 14:**
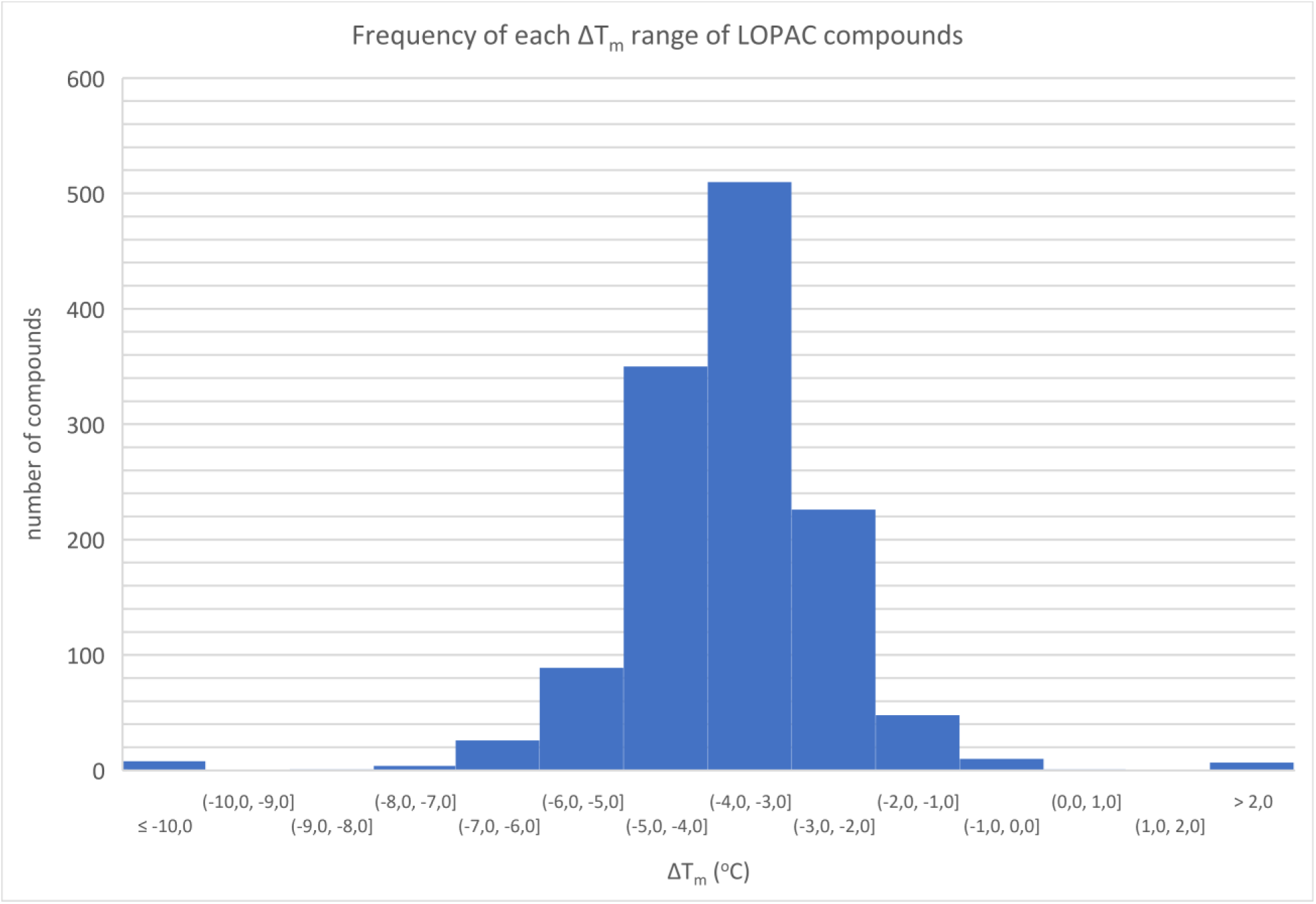
Histogram presenting the distribution of ΔT_m_ caused by LOPAC® compounds addition to SOD1(A4V). The ranges of ΔT_m_ are corresponded to the number of compounds that caused this T_m_ shift.

In order to select for target compounds to be further analyzed, some parameters should be established. For this selection, the readout is only the difference in T_m_ that each compound induces and not other features e.g. the alteration in the fluorescence (Matis et al., 2017; Münch & Bertolotti, 2010). Furthermore, several compounds induced a second shift, which is translated in the presence of a second T_m_ value upon their interaction with SOD1(A4V) and interestingly all cases shared the same pattern: the first shift was almost the same as SOD1(A4V) (protein as is) and the shift was located several degrees away. The second shift was clear and distinct as it was characterized by a clear increase in fluorescence which subsequently dropped and exhibited a minimum of the first derivative of its curve. The appearance of multiple transitions generally, can be attributed to different behavior of protein’s domains, aggregation, or ligands stabilizing a portion of the protein (Gao et al., 2020). In our case and since the first T_m_ is almost identical to the protein’s T_m_, it can be speculated, that this phenomenon occurs due to the chosen concentration of the ligand. Probably, a higher concentration would be adequate to interact with all molecules of the protein, which in that case would not leave molecules of SOD1(A4V) unbound to generate the initial protein’s T_m_.

### Selection of eleven compounds for their biochemical assessment as potent rescuers against SOD1(A4V) misfolding and aggregation

Initially, our interest was turned to the compounds presenting the biggest difference in ΔT_m_, both positive and negative. Concerning the positive differences, seven compounds (Diacylglycerol Kinase Inhibitor II, Cephalosporin C zinc salt, Cyclosporin A, Rabeprazole sodium, Calcimycin, Bexarotene and Icaritin) exhibited a positive ΔT_m_ ranging from 6°C till 32°C. As far as negative shifts are concerned, eight compounds (Mifamurtide, Carvedilol, CID11210285 hydrochloride, Idarubicin, Olvanil, N-Oleoylethanolamine, Dihydrocapsaicin, Artemether) presented ΔT_m_ ranging from - 10°C till −28°C. No matter how promising these large differences may seem, care should be taken for their interpretation. For this reason, the selection of the compounds is not only based on the ΔT_m_, but also the DSF spectra is taken into consideration. Since, ΔT_m_ is calculated based on the overall minimum of the first derivative of the curve (fluorescence vs temperature) the overview of the spectra is necessary to avoid false positive hits deriving from artifacts or mathematic errors upon derivatization. In addition, during library screens, sometimes it is the aggregation between molecules of the compounds themselves that generate leading to false results (Feng et al., 2007). Although, this problem is inevitable in most cases, it is resolved by further validation experiments.

It should be noted that a large shift of ΔT_m_ is not always correlated proportionally to the concentration or affinity of the ligands; larger shift in T_m_ does not imply tighter binding (Li, 2020; Sorrell et al., 2010; Vedadi et al., 2006; Waldron & Murphy, 2003). Different ligands interacting with the same protein bearing equivalent affinities, can result in different ΔT_m_. This phenomenon is related to the mechanism that each ligand uses to bind onto the protein; for instance, whether these interactions are more enthalpically or entropically driven. In case of large shifts, this type of interactions often leads to more entropically dominated interactions (McMahon et al., 2013).

As far as the decrease on the T_m_, it is possible to occur upon a drug binding on the protein. This can imply a non-specific binding or other mechanisms such as change in protein dynamics upon ligand binding (Li, 2020). Indeed ligands can destabilize proteins by binding primarily to their unfolded state and thus reduce the protein T_m_ (Cimmperman et al., 2008; Garbett & Chaires, 2012). Nevertheless, the negative shift does not exclude interaction and binding to the native state (Gao et al., 2020). Ligand binding to the unfolded protein state is a not well understood procedure since there are no data of crystal structures of any unfolded proteins to help our understanding (Cimmperman et al., 2008).

**Table 1:**
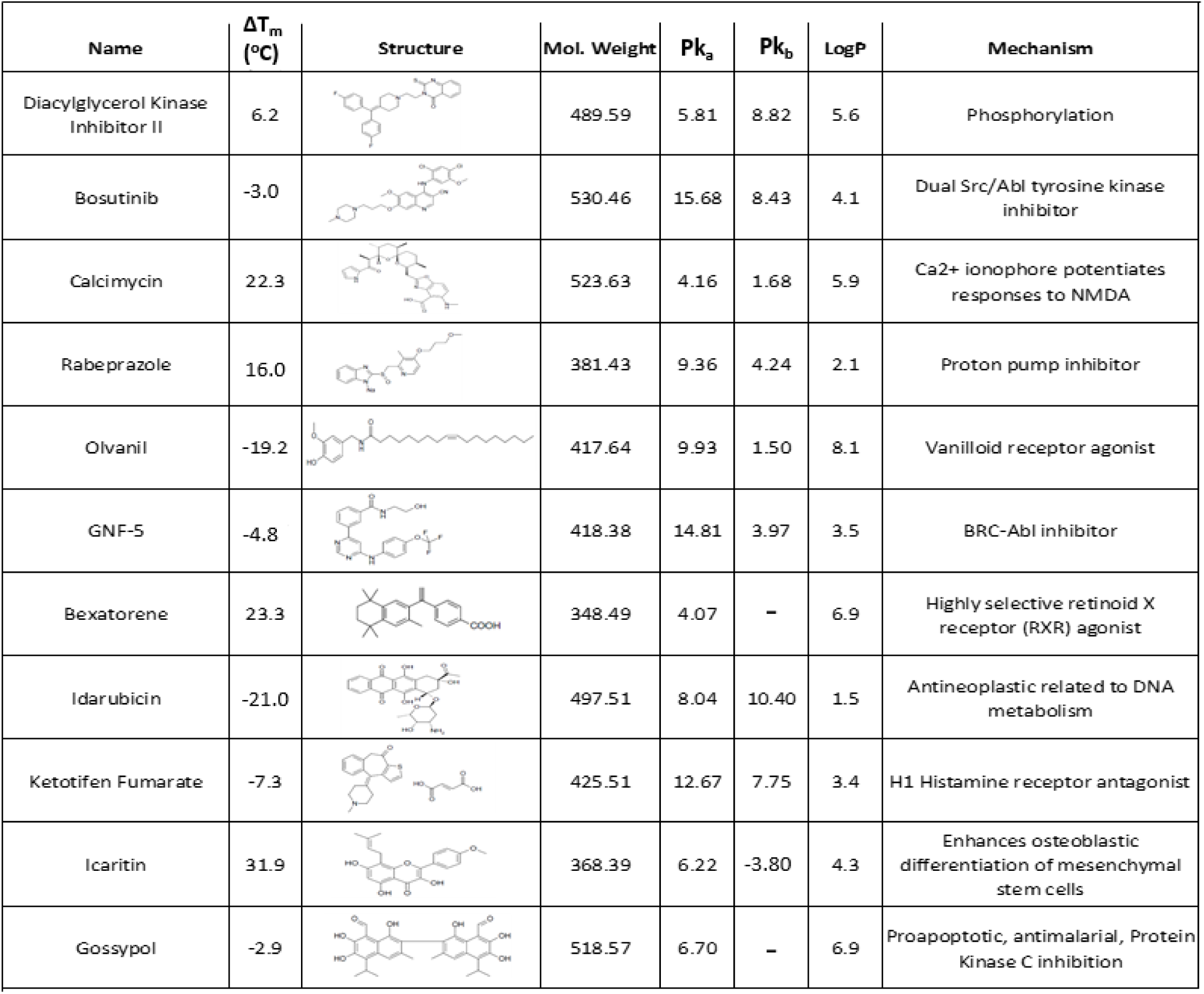
Lopac® compounds assessed to cause a significant ΔT_m_ on SOD1(A4V) and are worthy to be further examined. In this table some physicochemical, PK properties and their mechanism of actions are also presented.

| Name | $\Delta T_m$<br>(°C) | Structure | Mol. Weight | $Pk_a$ | $Pk_b$ | LogP | Mechanism |
| --- | --- | --- | --- | --- | --- | --- | --- |
| Diacylglycerol Kinase Inhibitor II | 6.2 |  | 489.59 | 5.81 | 8.82 | 5.6 | Phosphorylation |
| Bosutinib | -3.0 |  | 530.46 | 15.68 | 8.43 | 4.1 | Dual Src/Abl tyrosine kinase inhibitor |
| Calcimycin | 22.3 |  | 523.63 | 4.16 | 1.68 | 5.9 | Ca<sup>2+</sup> ionophore potentiates responses to NMDA |
| Rabeprazole | 16.0 |  | 381.43 | 9.36 | 4.24 | 2.1 | Proton pump inhibitor |
| Olvanil | -19.2 |  | 417.64 | 9.93 | 1.50 | 8.1 | Vanilloid receptor agonist |
| GNF-5 | -4.8 |  | 418.38 | 14.81 | 3.97 | 3.5 | BRC-Abl inhibitor |
| Bexatorene | 23.3 |  | 348.49 | 4.07 | - | 6.9 | Highly selective retinoid X receptor (RXR) agonist |
| Idarubicin | -21.0 |  | 497.51 | 8.04 | 10.40 | 1.5 | Antineoplastic related to DNA metabolism |
| Ketotifen Fumarate | -7.3 |  | 425.51 | 12.67 | 7.75 | 3.4 | H1 Histamine receptor antagonist |
| Icaritin | 31.9 |  | 368.39 | 6.22 | -3.80 | 4.3 | Enhances osteoblastic differentiation of mesenchymal stem cells |
| Gossypol | -2.9 |  | 518.57 | 6.70 | - | 6.9 | Proapoptotic, antimalarial, Protein Kinase C inhibition |

## DISCUSSION

SOD1 as holo-enzyme is one of the most thermostable proteins and thus theoretically would be expected to be the least, likely candidate protein molecule to undergo problematic folding (Furukawa et al., 2016).

However, presence of mutations endows the protein with the property of the tendency to misfold and aggregate. For the mutation analysed in this study, A4V, it not clear if the it has an effect on metal content of the protein: some published reports claim that this mutation is metallated *in vivo* (Elam et al., 2003) while others report that copper loading specifically drops significantly (Münch & Bertolotti, 2010; Ray et al., 2004). Based on our results, it seems that this mutation does not cause significant loss of enzyme’s activity.

We decided to employ a biophysical technique for the scanning of a chemical Library. This technique is DSF and is characterized by several advantages including the simplicity, the low cost of the reagents and its ability to perform medium-high throughput level of experiments. Despite that this type of studies examine interactions in an artificial environment that is void of other interactions and usually promotes misfolding and aggregation of the target protein, these cell-free platforms are a useful tool for choosing the compounds that will proceed to the next level of their evaluation including *in vivo* or animal studies.

We decided also to include this application in the contex of Drug repurposing (also known as drug repositioning or therapeutic switching), since, it is more advantageous in comparison to the classic drug discovery process due to the established safety profile of the drug, the shorter duration of the drug development phase and, accordingly the reduced cost along with the limited the risk of failure (Pushpakom et al., 2019; Shah et al., 2021). These features combined with the concomitant cost reduction it includes, creates a major advantage especially for ‘‘orphan diseases’’ like ALS (Pushpakom et al., 2019; Shah et al., 2021).

Interestingly, within the LOPAC® library were found compounds that have been associated with ALS disease in the past as possible disease modifiers and some of them such as ceftriaxone (C5793), bosutinib (PZ0192), minocycline (M9511), GABApentine (G154), cyclosporine (C3662), verapamil (V4629), celecoxib (PZ0008), carnitine (P4509), valproic acid (P4543), ropinirole (R2530), thalidomide (T144) have been tested in clinical trials II/III with inconclusive results or failure, that have shown promising results in ALS transgenic mice studies. Some others such as pentoxiffyline (P1784), nimodipine (N149), indinavir (SML0189) have also been tested in clinical trials ending up in not significant improvement on patients or failure but no data of transgenic SOD1 animal models were found in the literature meaning that they were chosen in the context of other mechanism like antioxidants. Finally, other compounds such as cisplatin (P4394), fluororacil (C1494) and isoproterenol (I2760) have a rescuing effect on SOD1 misfolding or stabilizing effect from *in vitro* studies in literature.

## AKNOWLEDGMENTS

**This research is co-financed by Greece and the European Union (European Social Fund-ESF) through the Operational Programme «Human Resources Development, Education and Lifelong Learning» in the context of the project “Strengthening Human Resources Research Potential via Doctorate Research – 2^nd^ Cycle” (MIS-5000432), implemented by the State Scholarships Foundation (IKY).**

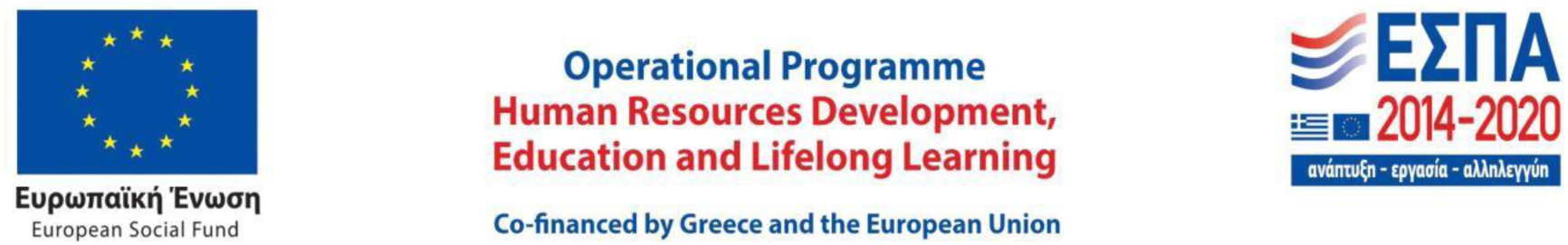

We would also like to thank Dr. Skretas Yiorgos for the kind gift of pASK75SOD1WT and pASK75SOD1(A4V) plasmids used for the production of human SOD1 recombinant protein.

