## Supplemental Table 1 for "Discovery and characterization of novel inhibitors against ALS-related SOD1(A4V) aggregation through screening of a chemical library using Differerential Scanning Fluorimetry (DSF)"

### ASSOCIATED CONTENT/ SUPPORTING INFORMATION

#### LOPAC® screening

**Table 1:** Table summarizing all the compounds from LOPAC® library and the difference in Tm of SOD1(A4V) they provoked. Compounds are presented by their catalogue number and the $\Delta T_m$ corresponds to the function: $T_m$ (SOD1(A4V))-$T_m$(SOD1(A4V) upon compound's presence). These results are presented according the value of the $\Delta T_m$ they caused; from bigger $T_m$ differences to smaller.

| Cat No | $\Delta T_m$ | Cat No | $\Delta T_m$ | Cat No | $\Delta T_m$ | Cat No | $\Delta T_m$ | Cat No | $\Delta T_m$ | Cat No | $\Delta T_m$ | Cat No | $\Delta T_m$ |
|---|---|---|---|---|---|---|---|---|---|---|---|---|---|
| SML0195 | -28.4 | **SML0666** | -4.7 | **T2265** | -4.1 | **S0752** | -3.8 | **R2751** | -3.4 | **A2731** | -2.9 | **S7198** | -2.2 |
| C3993 | -24.3 | **C1335** | -4.7 | **T5515** | -4.1 | **S7771** | -3.8 | **R5010** | -3.4 | **B1686** | -2.9 | **SML0744** | -2.2 |
| SML0698 | -21.2 | **C9847** | -4.7 | **T7508** | -4.1 | **SML0597** | -3.8 | **S0278** | -3.4 | **C-106** | -2.9 | **S2250** | -2.2 |
| I1656 | -21.0 | **D3648** | -4.7 | **U-100** | -4.1 | **B2515** | -3.7 | **S4063** | -3.4 | **D7802** | -2.9 | **B5002** | -2.1 |
| O0257 | -19.2 | **D4434** | -4.7 | **SML0584** | -4.1 | **B4558** | -3.7 | **S7395** | -3.4 | **E1279** | -2.9 | **D9071** | -2.1 |
| O0383 | -14.2 | **H2138** | -4.7 | **A2251** | -4.1 | **C0768** | -3.7 | **S8010** | -3.4 | **I6138** | -2.9 | **F4381** | -2.1 |
| M1022 | -10.9 | **I0782** | -4.7 | **A2385** | -4.1 | **C2505** | -3.7 | **B7148** | -3.3 | **K1003** | -2.9 | **U-116** | -2.1 |
| A9361 | -10.3 | **M1195** | -4.7 | **B5556** | -4.1 | **A-140** | -3.7 | **P5749** | -3.3 | **K1888** | -2.9 | **B-173** | -2.0 |
| P9375 | -8.7 | **C3909** | -4.7 | **B9061** | -4.1 | **A7275** | -3.7 | **PZ0129** | -3.3 | **L-134** | -2.9 | **E-114** | -2.0 |
| O2139 | -8.0 | **C4418** | -4.7 | **C4382** | -4.1 | **A7655** | -3.7 | **PZ0172** | -3.3 | **M3778** | -2.9 | **E7138** | -2.0 |
| 190047 | -7.9 | **C4520** | -4.7 | **C5040** | -4.1 | **A8423** | -3.7 | **SML0530** | -3.3 | **M4008** | -2.9 | **F3764** | -2.0 |
| K2628 | -7.3 | **C4662** | -4.7 | **C5259** | -4.1 | **A8598** | -3.7 | **A8981** | -3.3 | **N3288** | -2.9 | **I2892** | -2.0 |
| R-115 | -7.2 | **C5270** | -4.7 | **C8145** | -4.1 | **A9950** | -3.7 | **B7273** | -3.3 | **PZ0140** | -2.9 | **SML0040** | -2.0 |
| 265128 | -7.0 | **C7861** | -4.7 | **P-178** | -4.1 | **B6436** | -3.7 | **C1493** | -3.3 | **T-103** | -2.9 | **T9033** | -2.0 |
| A0606 | -6.9 | **C8395** | -4.7 | **P-215** | -4.1 | **D-103** | -3.7 | **D-031** | -3.3 | **T-144** | -2.9 | **L8668** | -2.0 |
| H5257 | -6.9 | **C8903** | -4.7 | **P7340** | -4.1 | **D-131** | -3.7 | **D-122** | -3.3 | **T2879** | -2.9 | **N2001** | -2.0 |
| A6566 | -6.7 | **S7701** | -4.6 | **P8688** | -4.1 | **D-193** | -3.7 | **D-134** | -3.3 | **T6050** | -2.9 | **N5504** | -2.0 |
| A7162 | -6.5 | **A5922** | -4.6 | **P9297** | -4.1 | **D9128** | -3.7 | **D14204** | -3.3 | **G8761** | -2.9 | **N5636** | -2.0 |
| 246379 | -6.5 | **A-201** | -4.6 | **P9797** | -4.1 | **E3256** | -3.7 | **D7910** | -3.3 | **N7510** | -2.9 | **P0453** | -2.0 |
| 246557 | -6.5 | **A-230** | -4.6 | **PZ0210** | -4.1 | **F6886** | -3.7 | **E0516** | -3.3 | **P2607** | -2.9 | **P0884** | -2.0 |
| A3085 | -6.5 | **A-254** | -4.6 | **Q0125** | -4.1 | **K4144** | -3.7 | **E2387** | -3.3 | **P4543** | -2.9 | **P4394** | -2.0 |

| | | | | | | | | | | | | | |
|---|---|---|---|---|---|---|---|---|---|---|---|---|---|
| A5626 | -6.4 | **A-255** | -4.6 | **S0693** | -4.1 | **L6668** | -3.7 | **E3132** | -3.3 | **C4024** | -2.9 | **SML0234** | -2.0 |
| I-119 | -6.4 | **A-263** | -4.6 | **S2816** | -4.1 | **PZ0014** | -3.7 | **E3876** | -3.3 | **C6506** | -2.9 | **E-006** | -1.9 |
| M2547 | -6.4 | **A8723** | -4.6 | **SML0524** | -4.1 | **S-009** | -3.7 | **F8927** | -3.3 | **C9758** | -2.9 | **F6300** | -1.9 |
| A0430 | -6.4 | **A9501** | -4.6 | **SML0704** | -4.1 | **S0568** | -3.7 | **N2288** | -3.3 | **P7791** | -2.9 | **PZ0196** | -1.9 |
| O7639 | -6.4 | **C1240** | -4.6 | **B-112** | -4.0 | **S-106** | -3.7 | **X1251** | -3.3 | **PZ0113** | -2.9 | **L9793** | -1.9 |
| PZ0162 | -6.4 | **F-131** | -4.6 | **B2390** | -4.0 | **SML0134** | -3.7 | **C-199** | -3.3 | **Q-110** | -2.9 | **N1016** | -1.9 |
| H2380 | -6.3 | **G2128** | -4.6 | **B-5311** | -4.0 | **T-123** | -3.7 | **C9911** | -3.3 | **R-108** | -2.9 | **N2034** | -1.9 |
| 861804 | -6.3 | **H-127** | -4.6 | **B7005** | -4.0 | **T4376** | -3.7 | **D0670** | -3.3 | **SML0752** | -2.9 | **N3136** | -1.9 |
| PZ0001 | -6.3 | **PZ0022** | -4.6 | **B7777** | -4.0 | **T6376** | -3.7 | **D1916** | -3.3 | **T7080** | -2.9 | **P1061** | -1.9 |
| A5376 | -6.2 | **S-168** | -4.6 | **B8385** | -4.0 | **T7313** | -3.7 | **D3630** | -3.3 | **C1610** | -2.8 | **T0826** | -1.9 |
| PZ0020 | -6.2 | **T1698** | -4.6 | **C0330** | -4.0 | **T8516** | -3.7 | **L8401** | -3.3 | **PZ0171** | -2.8 | **L3169** | -1.9 |
| A0382 | -6.2 | **T2896** | -4.6 | **C1671** | -4.0 | **U-101** | -3.7 | **M6545** | -3.3 | **C0256** | -2.8 | **SML0720** | -1.9 |
| D5446 | -6.2 | **T7540** | -4.6 | **S7067** | -4.0 | **W-104** | -3.7 | **M7445** | -3.3 | **D9305** | -2.8 | **B-121** | -1.8 |
| T3955 | -6.1 | **D3775** | -4.6 | **SML0564** | -4.0 | **Z0878** | -3.7 | **PZ0104** | -3.3 | **P0115** | -2.8 | **M-226** | -1.8 |
| A-265 | -6.1 | **D4526** | -4.6 | **A-022** | -4.0 | **Z2001** | -3.7 | **T0202** | -3.3 | **SML0218** | -2.8 | **F0778** | -1.7 |
| A4393 | -6.0 | **A0384** | -4.6 | **A-145** | -4.0 | **D1413** | -3.7 | **M-110** | -3.3 | **SML0229** | -2.8 | **F6145** | -1.7 |
| B7283 | -6.0 | **C3912** | -4.6 | **A8456** | -4.0 | **D1507** | -3.7 | **M-129** | -3.3 | **T7883** | -2.8 | **G6649** | -1.7 |
| M152 | -6.0 | **C4397** | -4.6 | **B0753** | -4.0 | **G5793** | -3.7 | **N-151** | -3.3 | **T9778** | -2.8 | **B6938** | -1.6 |
| P7912 | -6.0 | **C5020** | -4.6 | **B9311** | -4.0 | **H1877** | -3.7 | **N4148** | -3.3 | **U7500** | -2.8 | **E8875** | -1.6 |
| A4910 | -5.9 | **C6019** | -4.6 | **C-125** | -4.0 | **I4883** | -3.7 | **O2378** | -3.3 | **V8261** | -2.8 | **SML0536** | -1.6 |
| T9034 | -5.9 | **C7041** | -4.6 | **D5689** | -4.0 | **I9890** | -3.7 | **O3011** | -3.3 | **W1628** | -2.8 | **T8543** | -1.6 |
| A9561 | -5.9 | **O111** | -4.6 | **D5891** | -4.0 | **L-109** | -3.7 | **P2016** | -3.3 | **Y-102** | -2.8 | **A0233** | -1.6 |
| T3146 | -5.9 | **SML0221** | -4.6 | **D8296** | -4.0 | **L-121** | -3.7 | **PZ0213** | -3.3 | **B9685** | -2.8 | **A6605** | -1.6 |
| P8765 | -5.9 | **B3023** | -4.5 | **G-117** | -4.0 | **L9908** | -3.7 | **B4311** | -3.3 | **C-271** | -2.8 | **A5879** | -1.5 |
| D6518 | -5.8 | **B3501** | -4.5 | **G-154** | -4.0 | **M4659** | -3.7 | **P-120** | -3.3 | **D3900** | -2.8 | **A6134** | -1.5 |
| 291552 | -5.8 | **B5681** | -4.5 | **H0879** | -4.0 | **M6690** | -3.7 | **P-162** | -3.3 | **D5294** | -2.8 | **A6733** | -1.5 |
| A7148 | -5.7 | **C1625** | -4.5 | **H-135** | -4.0 | **PZ0100** | -3.7 | **P8293** | -3.3 | **L-135** | -2.8 | **N0630** | -1.5 |
| A-023 | -5.7 | **D-047** | -4.5 | **H5752** | -4.0 | **SML0264** | -3.7 | **P8828** | -3.3 | **S9311** | -2.8 | **P0547** | -1.5 |
| B8312 | -5.7 | **D3768** | -4.5 | **H9882** | -4.0 | **SML0667** | -3.7 | **P8887** | -3.3 | **SML0247** | -2.8 | **M7033** | -1.4 |
| D3689 | -5.7 | **F6889** | -4.5 | **I7379** | -4.0 | **U6881** | -3.7 | **P8891** | -3.3 | **C4915** | -2.8 | **R0529** | -1.4 |

| | | | | | | | | | | | | | |
|---|---|---|---|---|---|---|---|---|---|---|---|---|---|
| D5564 | -5.7 | **G3126** | -4.5 | **I9531** | -4.0 | **C3130** | -3.7 | **P9178** | -3.3 | **E5406** | -2.8 | **N2255** | -1.4 |
| A3134 | -5.7 | **H-168** | -4.5 | **L2536** | -4.0 | **C3412** | -3.7 | **R6250** | -3.3 | **N4159** | -2.8 | **P1675** | -1.4 |
| E-100 | -5.6 | **I7388** | -4.5 | **L3791** | -4.0 | **C8088** | -3.7 | **S0441** | -3.3 | **O3636** | -2.8 | **R1402** | -1.4 |
| G-119 | -5.6 | **M1275** | -4.5 | **L4762** | -4.0 | **C8138** | -3.7 | **S8251** | -3.3 | **P2116** | -2.8 | **R8875** | -1.4 |
| SML0527 | -5.6 | **T4512** | -4.5 | **L5025** | -4.0 | **C9510** | -3.7 | **B6311** | -3.2 | **R7772** | -2.8 | **A5006** | -1.3 |
| T6951 | -5.6 | **A0760** | -4.5 | **L9756** | -4.0 | **C9754** | -3.7 | **R1283** | -3.2 | **A0779** | -2.8 | **I7378** | -1.3 |
| C5793 | -5.6 | **A1260** | -4.5 | **M1809** | -4.0 | **E2535** | -3.7 | **A-236** | -3.2 | **R-116** | -2.8 | **M0814** | -1.3 |
| C8011 | -5.6 | **G5918** | -4.5 | **M5171** | -4.0 | **M-184** | -3.7 | **A4233** | -3.2 | **R-134** | -2.8 | **H9415** | -1.2 |
| A6351 | -5.5 | **N8652** | -4.5 | **M5435** | -4.0 | **O0766** | -3.7 | **A5181** | -3.2 | **R3255** | -2.8 | **E7881** | -1.1 |
| SML0505 | -5.5 | **P5514** | -4.5 | **M5560** | -4.0 | **P0878** | -3.7 | **D-030** | -3.2 | **S1438** | -2.8 | **D8941** | -1.0 |
| I1149 | -5.5 | **P7136** | -4.5 | **M5685** | -4.0 | **S3442** | -3.7 | **D-052** | -3.2 | **S1875** | -2.8 | **L-106** | -1.0 |
| SML0113 | -5.5 | **A3940** | -4.4 | **M7277** | -4.0 | **A0500** | -3.7 | **D-054** | -3.2 | **S8502** | -2.8 | **A9480** | -0.8 |
| T7165 | -5.5 | **A5330** | -4.4 | **PZ0003** | -4.0 | **I0658** | -3.7 | **E0137** | -3.2 | **PZ0185** | -2.7 | **I2285** | -0.8 |
| A1895 | -5.5 | **A8001** | -4.4 | **PZ0015** | -4.0 | **P-102** | -3.7 | **N1415** | -3.2 | **A7111** | -2.7 | **H8653** | -0.6 |
| A1910 | -5.5 | **B-169** | -4.4 | **PZ0139** | -4.0 | **P8782** | -3.7 | **N3911** | -3.2 | **B7688** | -2.7 | **N3510** | -0.6 |
| A5282 | -5.4 | **B8279** | -4.4 | **S-174** | -4.0 | **PZ0110** | -3.7 | **S-153** | -3.2 | **D6140** | -2.7 | **P1793** | -0.6 |
| A5909 | -5.4 | **S9318** | -4.4 | **SML0517** | -4.0 | **Q1250** | -3.7 | **S5317** | -3.2 | **D8555** | -2.7 | **S4250** | -0.6 |
| D9628 | -5.4 | **E4642** | -4.4 | **SML0634** | -4.0 | **Q3504** | -3.7 | **T0254** | -3.2 | **E4378** | -2.7 | **G6423** | -0.1 |
| H7779 | -5.4 | **F9552** | -4.4 | **SML0644** | -4.0 | **S7389** | -3.7 | **T-200** | -3.2 | **PZ0115** | -2.7 | **I8898** | 0.0 |
| I-120 | -5.4 | **G2536** | -4.4 | **SML0841** | -4.0 | **S8139** | -3.7 | **T7947** | -3.2 | **SML0075** | -2.7 | **SML0245** | 0.0 |
| P-152 | -5.4 | **G9797** | -4.4 | **T0410** | -4.0 | **B2377** | -3.6 | **W2270** | -3.2 | **SML0601** | -2.7 | **T2705** | 0.0 |
| SML0255 | -5.4 | **A-202** | -4.4 | **T1694** | -4.0 | **B6506** | -3.6 | **C5493** | -3.2 | **SML0892** | -2.7 | **SML0209** | 0.0 |
| SML0658 | -5.4 | **A7762** | -4.4 | **T6394** | -4.0 | **B7651** | -3.6 | **H7250** | -3.2 | **T2067** | -2.7 | **D5794** | 6.2 |
| M8046 | -5.4 | **A7845** | -4.4 | **T7040** | -4.0 | **C2755** | -3.6 | **I1637** | -3.2 | **T2408** | -2.7 | **C3270** | 12.1 |
| N-144 | -5.4 | **A9345** | -4.4 | **V8879** | -4.0 | **A6011** | -3.6 | **I6504** | -3.2 | **T3757** | -2.7 | **C3662** | 14.8 |
| 144509 | -5.4 | **B5437** | -4.4 | **X6000** | -4.0 | **PZ0002** | -3.6 | **I8021** | -3.2 | **U-120** | -2.7 | **SML0476** | 16.0 |
| A1755 | -5.4 | **C-239** | -4.4 | **Z4902** | -4.0 | **A-013** | -3.6 | **L1011** | -3.2 | **U4125** | -2.7 | **C7522** | 22.3 |

| | | | | | | | | | | | | | |
|---|---|---|---|---|---|---|---|---|---|---|---|---|---|
| S5890 | -5.4 | **D-027** | -4.4 | **C4479** | -4.0 | **A-143** | -3.6 | **L-119** | -3.2 | **V-100** | -2.7 | **SML0282** | 23.3 |
| A6671 | -5.3 | **D1414** | -4.4 | **C4522** | -4.0 | **A-164** | -3.6 | **M1514** | -3.2 | **D0540** | -2.7 | **SML0551** | 31.9 |
| A9512 | -5.3 | **D2521** | -4.4 | **C4895** | -4.0 | **D8399** | -3.6 | **M5154** | -3.2 | **H1753** | -2.7 | | |
| F6800 | -5.3 | **D4505** | -4.4 | **C4911** | -4.0 | **D9035** | -3.6 | **M5250** | -3.2 | **S4572** | -2.7 | | |
| V8138 | -5.3 | **D5782** | -4.4 | **C5982** | -4.0 | **F1553** | -3.6 | **B2185** | -3.2 | **SML0586** | -2.7 | | |
| C0494 | -5.3 | **D5886** | -4.4 | **C6643** | -4.0 | **F6020** | -3.6 | **L4545** | -3.2 | **M9651** | -2.7 | | |
| C-145 | -5.3 | **F4429** | -4.4 | **C7255** | -4.0 | **F9397** | -3.6 | **M9020** | -3.2 | **N1530** | -2.7 | | |
| F8682 | -5.3 | **F7932** | -4.4 | **C7971** | -4.0 | **H-140** | -3.6 | **N-142** | -3.2 | **N-170** | -2.7 | | |
| H7258 | -5.3 | **H1384** | -4.4 | **P-103** | -4.0 | **M6383** | -3.6 | **N7505** | -3.2 | **N4163** | -2.7 | | |
| H8502 | -5.3 | **H4759** | -4.4 | **P63204** | -4.0 | **P0111** | -3.6 | **N9007** | -3.2 | **N7758** | -2.7 | | |
| H8876 | -5.3 | **I7627** | -4.4 | **P8477** | -4.0 | **PZ0136** | -3.6 | **P4532** | -3.2 | **P0778** | -2.7 | | |
| H9003 | -5.3 | **J4137** | -4.4 | **P9879** | -4.0 | **S-154** | -3.6 | **P6126** | -3.2 | **P4668** | -2.7 | | |
| M1404 | -5.3 | **M1387** | -4.4 | **Q0875** | -4.0 | **S-180** | -3.6 | **P6628** | -3.2 | **T5575** | -2.7 | | |
| A1784 | -5.3 | **M7320** | -4.4 | **SML0678** | -4.0 | **SML0653** | -3.6 | **SML0227** | -3.2 | **A0937** | -2.7 | | |
| A7127 | -5.2 | **P9248** | -4.4 | **A4147** | -3.9 | **T0891** | -3.6 | **SML0552** | -3.2 | **R-121** | -2.7 | | |
| PZ0004 | -5.2 | **S3572** | -4.4 | **B-161** | -3.9 | **T1516** | -3.6 | **SML0630** | -3.2 | **R7150** | -2.7 | | |
| A9899 | -5.2 | **U-111** | -4.4 | **B-168** | -3.9 | **T1633** | -3.6 | **A8231** | -3.2 | **R8900** | -2.7 | | |
| B1381 | -5.2 | **N1392** | -4.4 | **B2134** | -3.9 | **T4182** | -3.6 | **C1494** | -3.2 | **S7882** | -2.7 | | |
| E7906 | -5.2 | **P0667** | -4.4 | **B7880** | -3.9 | **W-108** | -3.6 | **P-101** | -3.2 | **S8442** | -2.7 | | |
| F4765 | -5.2 | **P6503** | -4.4 | **C0400** | -3.9 | **Z-101** | -3.6 | **R2625** | -3.2 | **T6580** | -2.7 | | |
| P0122 | -5.2 | **S6319** | -4.4 | **C0424** | -3.9 | **D1306** | -3.6 | **R3277** | -3.2 | **D9175** | -2.6 | | |
| S-143 | -5.2 | **A0152** | -4.4 | **PZ0158** | -3.9 | **C4042** | -3.6 | **S3378** | -3.2 | **D9446** | -2.6 | | |

| | | | | | | | | | | | |
|---|---|---|---|---|---|---|---|---|---|---|---|
| C-104 | -5.2 | **A0487** | -4.4 | **SML0492** | -3.9 | **C5923** | -3.6 | **B-016** | -3.1 | **I9532** | -2.6 |
| H0131 | -5.2 | **A0966** | -4.4 | **E-101** | -3.9 | **C6022** | -3.6 | **B-138** | -3.1 | **K1136** | -2.6 |
| D9891 | -5.1 | **C7897** | -4.4 | **A-242** | -3.9 | **C7632** | -3.6 | **B5683** | -3.1 | **K3375** | -2.6 |
| E8375 | -5.1 | **R0758** | -4.4 | **C1618** | -3.9 | **C8031** | -3.6 | **B9308** | -3.1 | **M0253** | -2.6 |
| G-133 | -5.1 | **A6883** | -4.3 | **C-191** | -3.9 | **D6321** | -3.6 | **SML0520** | -3.1 | **M7945** | -2.6 |
| H9002 | -5.1 | **B-135** | -4.3 | **D1064** | -3.9 | **D8696** | -3.6 | **A3773** | -3.1 | **O7389** | -2.6 |
| I-122 | -5.1 | **B5016** | -4.3 | **D5766** | -3.9 | **I5409** | -3.6 | **A9605** | -3.1 | **R8404** | -2.6 |
| M2727 | -5.1 | **PZ0142** | -4.3 | **D7644** | -3.9 | **M-149** | -3.6 | **D-002** | -3.1 | **S9186** | -2.6 |
| P0099 | -5.1 | **SML0189** | -4.3 | **D9571** | -3.9 | **O3752** | -3.6 | **D126608** | -3.1 | **SML0275** | -2.6 |
| C-130 | -5.1 | **A-178** | -4.3 | **E1779** | -3.9 | **P7412** | -3.6 | **D-129** | -3.1 | **T9025** | -2.6 |
| A0257 | -5.1 | **A7250** | -4.3 | **E9531** | -3.9 | **P7561** | -3.6 | **D7505** | -3.1 | **X4753** | -2.6 |
| B8262 | -5.0 | **A7824** | -4.3 | **G-017** | -3.9 | **PZ0116** | -3.6 | **D8008** | -3.1 | **C-192** | -2.6 |
| C0625 | -5.0 | **D7443** | -4.3 | **G3796** | -3.9 | **R-103** | -3.6 | **D9190** | -3.1 | **A9861** | -2.6 |
| C1172 | -5.0 | **E5156** | -4.3 | **G7795** | -3.9 | **R6152** | -3.6 | **D9766** | -3.1 | **I7160** | -2.6 |
| F3680 | -5.0 | **F-124** | -4.3 | **I-135** | -3.9 | **T6950** | -3.6 | **H0126** | -3.1 | **M3071** | -2.6 |
| A2729 | -5.0 | **G8134** | -4.3 | **I-146** | -3.9 | **A4669** | -3.5 | **H8627** | -3.1 | **N7261** | -2.6 |
| A8835 | -5.0 | **O5889** | -4.3 | **I3639** | -3.9 | **C0737** | -3.5 | **I6659** | -3.1 | **N7778** | -2.6 |
| C9901 | -5.0 | **C3118** | -4.3 | **L9787** | -3.9 | **C0987** | -3.5 | **I8250** | -3.1 | **O8757** | -2.6 |
| D-003 | -5.0 | **D0676** | -4.3 | **M2901** | -3.9 | **A6664** | -3.5 | **L0664** | -3.1 | **O9126** | -2.6 |
| D2531 | -5.0 | **D5439** | -4.3 | **M3127** | -3.9 | **A9013** | -3.5 | **L2411** | -3.1 | **P4509** | -2.6 |
| E3263 | -5.0 | **H1252** | -4.3 | **M3184** | -3.9 | **C-101** | -3.5 | **L5647** | -3.1 | **P4670** | -2.6 |

| | | | | | | | | | | | |
|---|---|---|---|---|---|---|---|---|---|---|---|
| F4646 | -5.0 | **H2775** | -4.3 | **M4145** | -3.9 | **C-102** | -3.5 | **M1559** | -3.1 | **P6402** | -2.6 |
| H9772 | -5.0 | **H6036** | -4.3 | **M5441** | -3.9 | **C-117** | -3.5 | **M4910** | -3.1 | **T7205** | -2.6 |
| I2760 | -5.0 | **I0157** | -4.3 | **M6760** | -3.9 | **C-147** | -3.5 | **M5391** | -3.1 | **T8703** | -2.6 |
| L2167 | -5.0 | **I-114** | -4.3 | **M7319** | -3.9 | **C-203** | -3.5 | **M6500** | -3.1 | **X3629** | -2.6 |
| P9623 | -5.0 | **I-117** | -4.3 | **M7684** | -3.9 | **C-237** | -3.5 | **S-008** | -3.1 | **C-197** | -2.5 |
| SML00 15 | -5.0 | **L6545** | -4.3 | **N0287** | -3.9 | **D0411** | -3.5 | **S4443** | -3.1 | **D3634** | -2.5 |
| T1132 | -5.0 | **M2525** | -4.3 | **P0618** | -3.9 | **D1262** | -3.5 | **SML0226** | -3.1 | **D4007** | -2.5 |
| A1824 | -5.0 | **M2537** | -4.3 | **PZ0107** | -3.9 | **D-138** | -3.5 | **SML0613** | -3.1 | **D6899** | -2.5 |
| C7291 | -5.0 | **PZ0013** | -4.3 | **PZ0114** | -3.9 | **D2926** | -3.5 | **U6758** | -3.1 | **E2031** | -2.5 |
| P6909 | -5.0 | **T1505** | -4.3 | **PZ0137** | -3.9 | **D7938** | -3.5 | **Y3125** | -3.1 | **E3380** | -2.5 |
| R0875 | -5.0 | **T4143** | -4.3 | **PZ0141** | -3.9 | **E0381** | -3.5 | **C-108** | -3.1 | **E4375** | -2.5 |
| A0232 | -4.9 | **T4425** | -4.3 | **S4528** | -3.9 | **E1896** | -3.5 | **D1260** | -3.1 | **S8688** | -2.5 |
| F2802 | -4.9 | **C7005** | -4.3 | **S5192** | -3.9 | **E2375** | -3.5 | **D0943** | -3.1 | **SML0610** | -2.5 |
| A9834 | -4.9 | **C8645** | -4.3 | **T1443** | -3.9 | **F6513** | -3.5 | **N3398** | -3.1 | **SML0824** | -2.5 |
| C6492 | -4.9 | **M5793** | -4.3 | **T6031** | -3.9 | **G0668** | -3.5 | **N3529** | -3.1 | **SML0864** | -2.5 |
| G-002 | -4.9 | **N8534** | -4.3 | **U-105** | -3.9 | **H1512** | -3.5 | **N7634** | -3.1 | **B5435** | -2.5 |
| G0419 | -4.9 | **P1918** | -4.3 | **U5882** | -3.9 | **I-138** | -3.5 | **O-100** | -3.1 | **M-001** | -2.5 |
| H0627 | -4.9 | **P5396** | -4.3 | **V4629** | -3.9 | **I5627** | -3.5 | **O2881** | -3.1 | **N-153** | -2.5 |
| H8125 | -4.9 | **PZ0111** | -4.3 | **V6383** | -3.9 | **L-131** | -3.5 | **O9637** | -3.1 | **N-183** | -2.5 |
| I0375 | -4.9 | **P-203** | -4.3 | **N8403** | -3.9 | **L4376** | -3.5 | **P-105** | -3.1 | **N4382** | -2.5 |
| I2279 | -4.9 | **P8813** | -4.3 | **N8659** | -3.9 | **L8533** | -3.5 | **P1784** | -3.1 | **P2278** | -2.5 |

| | | | | | | | | | | | |
|---|---|---|---|---|---|---|---|---|---|---|---|
| I4409 | -4.9 | **R5648** | -4.3 | **P5295** | -3.9 | **M1777** | -3.5 | **P1801** | -3.1 | **PZ0102** | -2.5 |
| L4900 | -4.9 | **B2417** | -4.2 | **P5654** | -3.9 | **M5644** | -3.5 | **P4651** | -3.1 | **R2530** | -2.5 |
| M6191 | -4.9 | **G6548** | -4.2 | **A1237** | -3.9 | **M6524** | -3.5 | **P6499** | -3.1 | **S2201** | -2.5 |
| SML0216 | -4.9 | **A8404** | -4.2 | **A1980** | -3.9 | **M6628** | -3.5 | **P7295** | -3.1 | **S3065** | -2.5 |
| SML0269 | -4.9 | **C9369** | -4.2 | **B5063** | -3.9 | **O1141** | -3.5 | **P8852** | -3.1 | **SML0683** | -2.5 |
| SML0660 | -4.9 | **D-108** | -4.2 | **P6656** | -3.9 | **PZ0016** | -3.5 | **PZ0101** | -3.1 | **A4562** | -2.4 |
| N5023 | -4.9 | **D5676** | -4.2 | **P6777** | -3.9 | **S-006** | -3.5 | **PZ0117** | -3.1 | **B-134** | -2.4 |
| P3510 | -4.9 | **D6035** | -4.2 | **P8013** | -3.9 | **SML0223** | -3.5 | **R0500** | -3.1 | **C0862** | -2.4 |
| 194336 | -4.9 | **F-114** | -4.2 | **P9233** | -3.9 | **T0625** | -3.5 | **S5321** | -3.1 | **F4303** | -2.4 |
| 211672 | -4.9 | **F9677** | -4.2 | **P9547** | -3.9 | **T-104** | -3.5 | **SML0711** | -3.1 | **F8257** | -2.4 |
| A3145 | -4.9 | **T2057** | -4.2 | **PZ0112** | -3.9 | **T6943** | -3.5 | **B-102** | -3.0 | **N2538** | -2.4 |
| C3930 | -4.9 | **V1377** | -4.2 | **R9525** | -3.9 | **V7264** | -3.5 | **C2538** | -3.0 | **PZ0012** | -2.4 |
| C6895 | -4.9 | **G-007** | -4.2 | **S2812** | -3.9 | **C8773** | -3.5 | **D3943** | -3.0 | **SML0639** | -2.4 |
| G5794 | -4.9 | **H8645** | -4.2 | **S7936** | -3.9 | **E9658** | -3.5 | **A1362** | -3.0 | **T5648** | -2.4 |
| P6902 | -4.9 | **H9876** | -4.2 | **SML0550** | -3.9 | **H8664** | -3.5 | **B6813** | -3.0 | **D2763** | -2.4 |
| Q-109 | -4.9 | **I-106** | -4.2 | **Z2777** | -3.9 | **P-118** | -3.5 | **C0996** | -3.0 | **H4001** | -2.4 |
| S7947 | -4.9 | **I7016** | -4.2 | **C2235** | -3.8 | **P7780** | -3.5 | **C1119** | -3.0 | **T-122** | -2.4 |
| A4638 | -4.8 | **M0763** | -4.2 | **A-155** | -3.8 | **P8511** | -3.5 | **C3618** | -3.0 | **Y0503** | -2.4 |
| B4555 | -4.8 | **M1818** | -4.2 | **A6476** | -3.8 | **PZ0108** | -3.5 | **D-104** | -3.0 | **M-104** | -2.4 |
| C1159 | -4.8 | **M2398** | -4.2 | **A8676** | -3.8 | **Q3251** | -3.5 | **F0881** | -3.0 | **N5501** | -2.4 |
| A-129 | -4.8 | **PZ0211** | -4.2 | **A9251** | -3.8 | **R-104** | -3.5 | **F-100** | -3.0 | **O0250** | -2.4 |

| | | | | | | | | | | | |
|---|---|---|---|---|---|---|---|---|---|---|---|
| A8003 | -4.8 | **SML0091** | -4.2 | **A9809** | -3.8 | **R7385** | -3.5 | **F1016** | -3.0 | **O1008** | -2.4 |
| A9256 | -4.8 | **SML0130** | -4.2 | **C8863** | -3.8 | **R9644** | -3.5 | **G6416** | -3.0 | **P4015** | -2.4 |
| B2009 | -4.8 | **SML0179** | -4.2 | **D-1920-6** | -3.8 | **S2876** | -3.5 | **I-160** | -3.0 | **SML0612** | -2.4 |
| E3149 | -4.8 | **C3635** | -4.2 | **D8040** | -3.8 | **S5013** | -3.5 | **I18008** | -3.0 | **R-140** | -2.4 |
| G0639 | -4.8 | **C4238** | -4.2 | **H3288** | -3.8 | **S8197** | -3.5 | **K0250** | -3.0 | **T2952** | -2.4 |
| R0158 | -4.8 | **C5134** | -4.2 | **PZ0178** | -3.8 | **A3539** | -3.4 | **K1751** | -3.0 | **Y4877** | -2.3 |
| S3316 | -4.8 | **C6645** | -4.2 | **SML0594** | -3.8 | **B2292** | -3.4 | **L0258** | -3.0 | **A8852** | -2.3 |
| SML0776 | -4.8 | **M-137** | -4.2 | **T9652** | -3.8 | **C1754** | -3.4 | **L4408** | -3.0 | **D6940** | -2.3 |
| T-101 | -4.8 | **N-115** | -4.2 | **C-141** | -3.8 | **S3322** | -3.4 | **M2776** | -3.0 | **F6627** | -2.3 |
| A7606 | -4.8 | **O3125** | -4.2 | **C-223** | -3.8 | **A8054** | -3.4 | **M3262** | -3.0 | **G4796** | -2.3 |
| H-108 | -4.8 | **P6500** | -4.2 | **D1542** | -3.8 | **C-144** | -3.4 | **M3281** | -3.0 | **PZ0008** | -2.3 |
| H1377 | -4.8 | **A2129** | -4.2 | **I2764** | -3.8 | **C-231** | -3.4 | **M3668** | -3.0 | **B175** | -2.3 |
| N1786 | -4.8 | **P7505** | -4.2 | **J4829** | -3.8 | **D5290** | -3.4 | **M3953** | -3.0 | **H8250** | -2.3 |
| T2528 | -4.8 | **P8139** | -4.2 | **L9664** | -3.8 | **D5385** | -3.4 | **M4796** | -3.0 | **I3766** | -2.3 |
| C6048 | -4.8 | **B2640** | -4.1 | **M7065** | -3.8 | **E1383** | -3.4 | **PZ0011** | -3.0 | **M-105** | -2.3 |
| C6628 | -4.8 | **B3650** | -4.1 | **SML0679** | -3.8 | **K3888** | -3.4 | **PZ0121** | -3.0 | **M-107** | -2.3 |
| C7912 | -4.8 | **B5275** | -4.1 | **SML0777** | -3.8 | **L2037** | -3.4 | **S3567** | -3.0 | **M9292** | -2.3 |
| I0160 | -4.8 | **B5399** | -4.1 | **T1512** | -3.8 | **L2906** | -3.4 | **S5567** | -3.0 | **M9656** | -2.3 |
| M9511 | -4.8 | **C0750** | -4.1 | **T4264** | -3.8 | **L5783** | -3.4 | **S8822** | -3.0 | **N-158** | -2.3 |
| PZ0135 | -4.8 | **C1251** | -4.1 | **T4500** | -3.8 | **M4531** | -3.4 | **SML0817** | -3.0 | **O0877** | -2.3 |
| 218359 | -4.8 | **C1290** | -4.1 | **T5576** | -3.8 | **M5379** | -3.4 | **T7665** | -3.0 | **PZ0124** | -2.3 |
| SML0511 | -4.8 | **A4508** | -4.1 | **T5625** | -3.8 | **M6517** | -3.4 | **U-103** | -3.0 | **SML0246** | -2.3 |

| | | | | | | | | | | | |
|---|---|---|---|---|---|---|---|---|---|---|---|
| A3711 | -4.7 | **A-138** | -4.1 | **T6764** | -3.8 | **SML0233** | -3.4 | **U-104** | -3.0 | **E9750** | -2.2 |
| A7009 | -4.7 | **A-162** | -4.1 | **C4542** | -3.8 | **T0318** | -3.4 | **U-106** | -3.0 | **F8791** | -2.2 |
| A7410 | -4.7 | **D7909** | -4.1 | **C7230** | -3.8 | **T6164** | -3.4 | **Y-101** | -3.0 | **SML0252** | -2.2 |
| A9657 | -4.7 | **D9815** | -4.1 | **L5420** | -3.8 | **T7254** | -3.4 | **C6305** | -3.0 | **T9262** | -2.2 |
| A9699 | -4.7 | **E4660** | -4.1 | **M2199** | -3.8 | **U-115** | -3.4 | **C8221** | -3.0 | **K3519** | -2.2 |
| B0385 | -4.7 | **F9427** | -4.1 | **P5052** | -3.8 | **Z1252** | -3.4 | **D-033** | -3.0 | **M-116** | -2.2 |
| B1266 | -4.7 | **G-110** | -4.1 | **120693** | -3.8 | **M-108** | -3.4 | **F0430** | -3.0 | **M4699** | -2.2 |
| C6617 | -4.7 | **H-128** | -4.1 | **A2169** | -3.8 | **M8131** | -3.4 | **K1015** | -3.0 | **M9125** | -2.2 |
| F-132 | -4.7 | **H9523** | -4.1 | **G9048** | -3.8 | **N-149** | -3.4 | **P9391** | -3.0 | **N-156** | -2.2 |
| G6793 | -4.7 | **I5879** | -4.1 | **P-126** | -3.8 | **N8784** | -3.4 | **PZ0192** | -3.0 | **N4779** | -2.2 |
| N2540 | -4.7 | **L8539** | -4.1 | **P-154** | -3.8 | **O0886** | -3.4 | **S0758** | -3.0 | **N5751** | -2.2 |
| PZ0005 | -4.7 | **P9872** | -4.1 | **P8386** | -3.8 | **P5679** | -3.4 | **S2501** | -3.0 | **N7127** | -2.2 |
| PZ0007 | -4.7 | **PZ0106** | -4.1 | **PZ0103** | -3.8 | **C3025** | -3.4 | **SML0518** | -3.0 | **O2751** | -2.2 |
| S2671 | -4.7 | **SML0191** | -4.1 | **Q-102** | -3.8 | **I5784** | -3.4 | **SML0557** | -3.0 | **P0130** | -2.2 |
| S8567 | -4.7 | **SML0210** | -4.1 | **Q2128** | -3.8 | **R-101** | -3.4 | **Z4626** | -3.0 | **P4405** | -2.2 |
